# Structural determination of *Rickettsia* lipid A without chemical extraction confirms shorter acyl chains in later-evolving Spotted Fever Group pathogens

**DOI:** 10.1101/2023.07.06.547954

**Authors:** Hyojik Yang, Victoria I. Verhoeve, Courtney E. Chandler, Shreeram Nallar, Greg A. Snyder, Robert K. Ernst, Joseph J. Gillespie

**Affiliations:** Department of Microbial Pathogenesis, School of Dentistry, University of Maryland, Baltimore, MD 21201; Department of Microbiology and Immunology, School of Medicine, University of Maryland Baltimore, MD 21201; Division of Vaccine Research, Institute of Human Virology, University of Maryland Baltimore, MD 21201

**Keywords:** *Rickettsia*, lipopolysaccharide, lipid A, FLAT^n^, pathogenesis, Rickettsiosis, Spotted Fever Group, evolution, Rocky Mountain Spotted Fever

## Abstract

Rickettsiae are Gram-negative obligate intracellular parasites of numerous eukaryotes. Human pathogens of the Transitional Group (TRG), Typhus Group (TG), and Spotted Fever Group (SFG) rickettsiae infect blood-feeding arthropods, have dissimilar clinical manifestations, and possess unique genomic and morphological attributes. Lacking glycolysis, rickettsiae pilfer numerous metabolites from host cytosol to synthesize peptidoglycan and lipopolysaccharide (LPS). For LPS, O-antigen immunogenicity varies between SFG and TG pathogens; however, lipid A proinflammatory potential is unknown. We previously demonstrated that *R. akari* (TRG), *R. typhi* (TG), and *R. montanensis* (SFG) produce lipid A with long 2’ secondary acyl chains (C16 or C18) compared to short 2’ secondary acyl chains (C12) in *R. rickettsii* (SFG) lipid A. To further probe this structural heterogeneity and estimate a time point when shorter 2’ secondary acyl chains originated, we generated lipid A structures for two additional SFG rickettsiae (*R. rhipicephali* and *R. parkeri*) utilizing Fast Lipid Analysis Technique adopted for use with tandem mass spectrometry (FLAT^n^). FLAT^n^ allowed analysis of lipid A structure directly from host cell-purified bacteria, providing substantial improvement over lipid A chemical extraction. FLAT^n^-derived structures indicate SFG rickettsiae diverging after *R. rhipicephali* evolved shorter 2’ secondary acyl chains. Bioinformatics analysis of *Rickettsia* LpxL late acyltransferases revealed discrete active sites and hydrocarbon rulers for long versus short 2’ secondary acyl chain addition. While the significance of different lipid A structures for diverse *Rickettsia* pathogens is unknown, our success using FLAT^n^ will facilitate determining how structural heterogeneity impacts interactions with host lipid A receptors and overall inflammatory potential.

**IMPORTANCE:** Deforestation, urbanization, and homelessness lead to spikes in Rickettsioses. Vector-borne human pathogens of Transitional Group (TRG), Typhus Group (TG), and Spotted Fever Group (SFG) rickettsiae differ by clinical manifestations, immunopathology, genome composition, and morphology. We previously showed that lipid A (or endotoxin), the membrane anchor of Gram-negative bacterial lipopolysaccharide (LPS), structurally differs in *R. rickettsii* (later-evolving SFG) relative to *R. montanensis* (basal SFG), *R. typhi* (TG), and *R. akari* (TRG). As lipid A structure influences recognition potential in vertebrate LPS sensors, further assessment of *Rickettsia* lipid A structural heterogeneity is needed. Here, we sidestepped the difficulty of *ex vivo* lipid A chemical extraction by utilizing FLAT^n^, a new procedure for generating lipid A structures directly from host cell-purified bacteria. These data confirm later-evolving SFG pathogens synthesize structurally distinct lipid A. Our findings impact interpreting immune responses to different *Rickettsia* pathogens and utilizing lipid A adjuvant or anti-inflammatory properties in vaccinology.

## OBSERVATION

Genus *Rickettsia* (*Alphaproteobacteria*: Rickettsiales) comprises species of obligate intracellular parasites of numerous eukaryotes (1). Across the *Rickettsia* phylogeny, all known agents of human disease are vector-borne pathogens of the **Transitional Group** (**TRG**), **Typhus Group** (**TG**), or **Spotted Fever Group** (**SFG**) rickettsiae, which are later-evolving clades relative to basal lineages of numerous invertebrate and protist endosymbionts (2). Compared to other Rickettsiales species with human health relevance (e.g., *Orientia tsutsugamushi*, wolbachiae, species of *Neorickettsia, Anaplasma, Neoehrlichia* and *Ehrlichia*), rickettsiae alone synthesize **lipopolysaccharide** (**LPS**) (3), which is comprised of extracellular polysaccharide chains (O-antigen) linked to a membrane phosphoglycolipid (lipid A) by a core oligosaccharide. As all *Rickettsia* species lack glycolytic enzymes, they are the only known bacteria to synthesize a canonical Gram-negative cell envelope rich in LPS, as well as peptidoglycan (4, 5), from metabolites derived from host cytosol (6).

All described rickettsioses are caused by arthropod-borne pathogens that differ in their protein secretome (7, 8) and presumably LPS composition. For LPS, *Rickettsia* O-antigen contains the sugar quinovosamine (9–11) that is required for S-layer formation and vertebrate pathogenicity (12). While the proinflammatory nature of *Rickettsia* lipid A remains unknown, it is a candidate for the well characterized triggering of mammalian MD-2/TLR4 receptor and non-canonical inflammasome activation during infection (13–18). We previously demonstrated that *R. akari* (TRG), *R. typhi* (TG), and *R. montanensis* (SFG) produce lipid A with longer 2’ secondary acyl chains (C16 or C18) as compared to shorter chains (C12) in *R. rickettsii* (SFG) lipid A (19). As *R. rickettsii* is later-evolving in the SFG rickettsiae relative to *R. montanensis*, we surmised that a switch from longer to shorter 2’ secondary acyl chains occurred later in SFG rickettsial evolution. Importantly, this later-evolving clade is dominated by other notable human pathogens, including the agents of Japanese spotted fever (*R. japonica*), Flinders Island spotted fever (*R. honei*), Pacific Coast tick fever (*R. philipii*), Mediterranean spotted fever (*R. conorii*), Siberian tick typhus (*R. sibirica*), African tick-bite fever (*R. africae*), and *Dermacentor*-borne necrosis erythema and lymphadenopathy (*R. raoultii* and *R. slovaca*).

To further evaluate *Rickettsia* lipid A structural heterogeneity, we selected two additional species, *R. rhipicephali* and *R. parkeri*, for lipid A structural analysis. In recent phylogeny estimations, *R. rhipicephali* groups in a clade that diverges after *R. montanensis* (long 2’ secondary acyl chains) but before *R. rickettsii* (short 2’ secondary acyl chain), while *R. parkeri* belongs to a clade that is sister to the *R. rickettsii*-containing clade (1, 7). Thus, our strategy zeros in on the evolutionary timepoint for the transition to short 2’ secondary acyl chains in SFG rickettsiae. Further, we utilized a different analytical approach to generate new structures called **Fast Lipid Analysis Technique adopted for use with tandem mass spectrometry** (**FLAT**^**n**^), allowing assessment of prior structure determinations that were based on lipid A chemical extraction. Briefly, FLAT^n^ is a method for on-surface and/or on-tissue release of lipid A from LPS that allows its detection by **matrix-assisted laser desorption ionization** (**MALDI**) **mass spectrometry** (**MS**) in the negative ion mode (20, 21) and has been shown to facilitate direct analysis of lipid A structure from a single bacteria colony (22). Thus, FLAT^n^ circumvents time-, labor-, and sample-intensive techniques previously required to chemically extract lipid A prior to MS analysis. Thus, we reasoned that FLAT^n^ would yield *Rickettsia* lipid A structures from a minimal sample of bacteria partially purified from far fewer host cells than our prior approach.

FLAT^n^ allowed for the direct analysis of *R. rhipicephali* and *R. parkeri* lipid A structures from host cells, providing substantial improvement and efficiency over lipid A chemical extraction (**FIG. 1**). Bacterial samples that were purified from hosts cells using either bead- or sucrose gradient-based strategies sufficed to generate MS spectra at either 1936.37 *m/z* (C16 2’ secondary acyl chains) or 1880.31 *m/z* (C12 2’ secondary acyl chains), consistent with prior analyses of chemically-extracted *Rickettsia* lipid A (19) (**FIG. 1A, B**). Subsequent derivatization of these single ions for *R. rhipicephali* (**FIG. 1C**) and *R. parkeri* (**FIG. 1D**) yielded fragmentation products that supported structural elucidation. The FLAT^n^-derived structure for *R. rhipicephali* lipid A corroborates those of *R. akari, R. typhi*, and *R. montanensis* with long 2’ secondary acyl chains (**FIG. 1E**). In contrast, the *R. parkeri* FLAT^n^-derived lipid A structure matches that determined for *R. rickettsii* strains with short 2’ secondary acyl chains (**FIG. 1F**). Thus, as with prior results (19), we have defined a trait in later-evolving SFG rickettsiae lipid A potentially leading to a more inflammatory lipid A structure observed in Enterobacteriaceae species.

**FIGURE 1.**
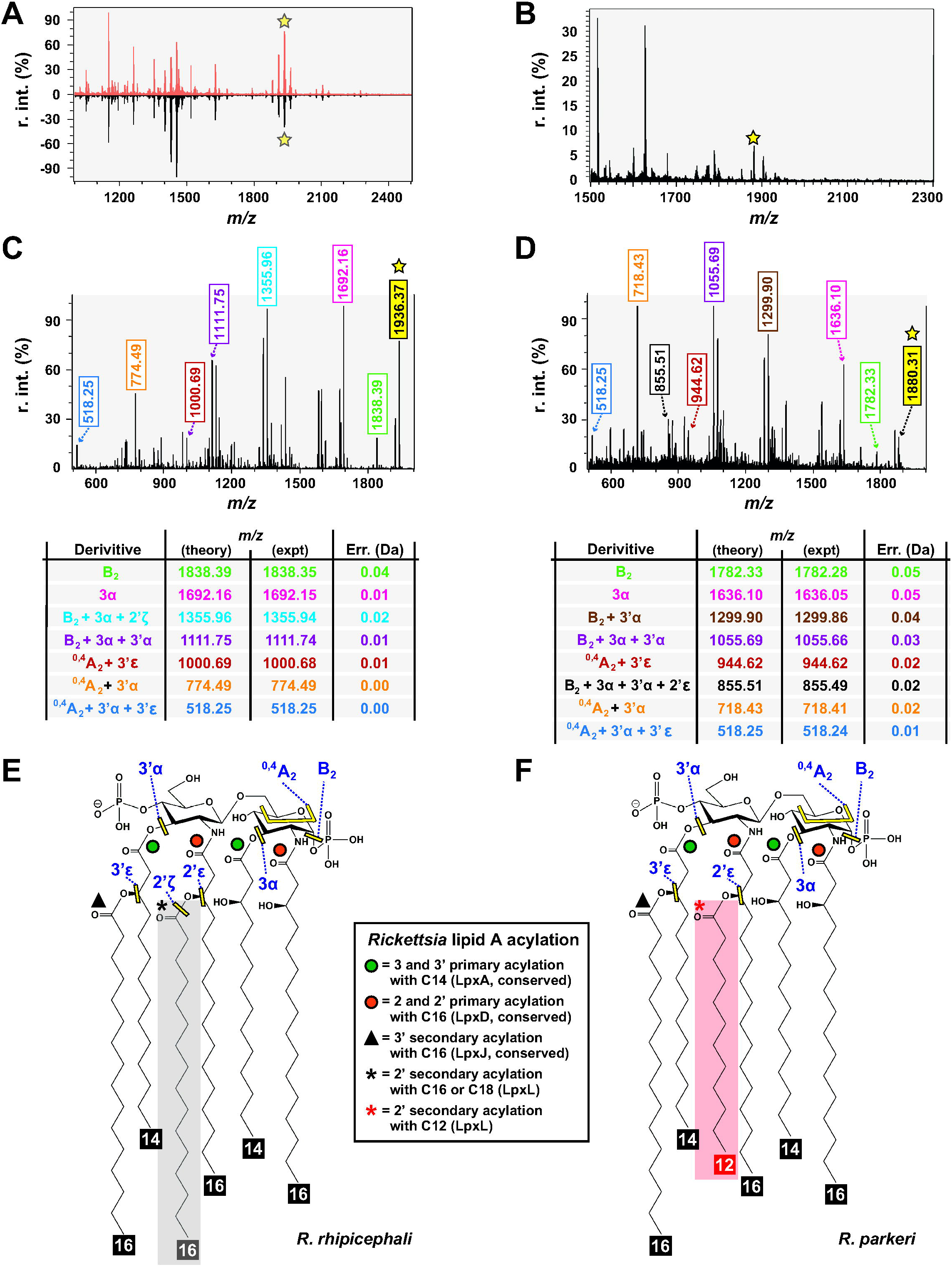
*Rickettsia* Lipid A structures determined by FLAT^n^ confirm variable acyl chain lengths for different rickettsiae. **(A, B)** FLAT-MS spectra from **(A)** *R. rhipicephali* partially purified from host cells using either a bead (B) or sucrose gradient (S) purification strategy, and from **(B)** *R. parkeri* partially purified from host cells using sucrose gradient purification strategy. Briefly, Vero76 cells were grown to confluence in T-25 flasks with cells infected at an MOI10. At 3 days post infection, host cells were recovered and lysed with 3mm beads, host debris removed via low speed centrifugation, and rickettsia collected by high speed centrifugation. Purification via sucrose gradient followed our prior protocol (19). Bacteria pellets were then analyzed via FLATn. Stars indicate the expected size for *Rickettsia* lipid A with C16 (∼1936.37 m/z) or C12 (1880.31 m/z) 2’ secondary acyl chains based on our prior report (19). **(C, D)** Derivatization of a single ion for the **(C)** *R. rhipicephali* sample (1936.37 m/z) and the **(D)** *R. parkeri* sample illustrating five and six, respectively, major fragmentation products. These products are named in the tables with theoretical and experimental sizes shown, with error calculation illustrating robust interpretation. They are also colored-coded to facilitate interpretation of the spectra above and predicted structures. **(E, F)** FLAT^n^-derived structure predictions for lipid A of **(E)** *R. rhipicephali*, which is similar to previously determined *R. akari, R. typhi* and *R. montanensis* structures, and **(F)** *R. parkerii*, which is similar to previously determined structures for *R. rickettsii* strains Shelia Smith and Iowa. Sites yielding fragmentation products are yellow, with corresponding nomenclature described in the tables in panels **C** and **D**. The inset describes the conserved and variable lipid A acylation of *Rickettsia* lipid A, with colored symbols mapped on structures (see **Fig. S1** for more details).

While bolstering prior results that suggested *Rickettsia* lipid A structural heterogeneity, FLAT^n^-derived structures also helped refine the evolutionary time point when short 2’ secondary acyl chains originated within SFG rickettsiae (**FIG. 2A**). In light of phylogeny estimation, our collective data indicate SFG rickettsiae diverging after *R. rhipicephali* evolved shorter 2’ secondary acyl chains (**FIG. 2A**, red shading). A parsimonious interpretation entails all members of the *R. rhipicephali*-containing clade synthesize lipid A with long 2’ secondary acyl chains, whereas all later-evolving SFG rickettsiae synthesize lipid A with short 2’ secondary acyl chains. Aside from many human pathogens in the later-evolving lineages, *R. peacockii* and the endosymbiont of the seal fur louse (*Proechinophthirus fluctus*) are non-pathogenic species that do not infect vertebrates (23, 24). Assuming that these endosymbionts synthesize lipid A (both of their genomes encode the full suite of Raetz pathway enzymes (3)), short 2’ secondary acyl chains could be considered a trait that emerged without selective pressure from vertebrate host environment. Alternatively, the non-pathogenic endosymbionts of this clade may have lost the ability to invade vertebrate hosts due to pseudogenization of numerous genes implicated in vertebrate cell invasion (23, 24), including uncharacterized LPS-modification enzymes that may only be necessary for survival in vertebrate hosts (25). Generating lipid A structures for other species in this clade will be necessary to further evaluate our observations, as will determining if rickettsiae alter their lipid A acyl chain lengths in vertebrate versus invertebrate hosts.

**FIGURE 2.**
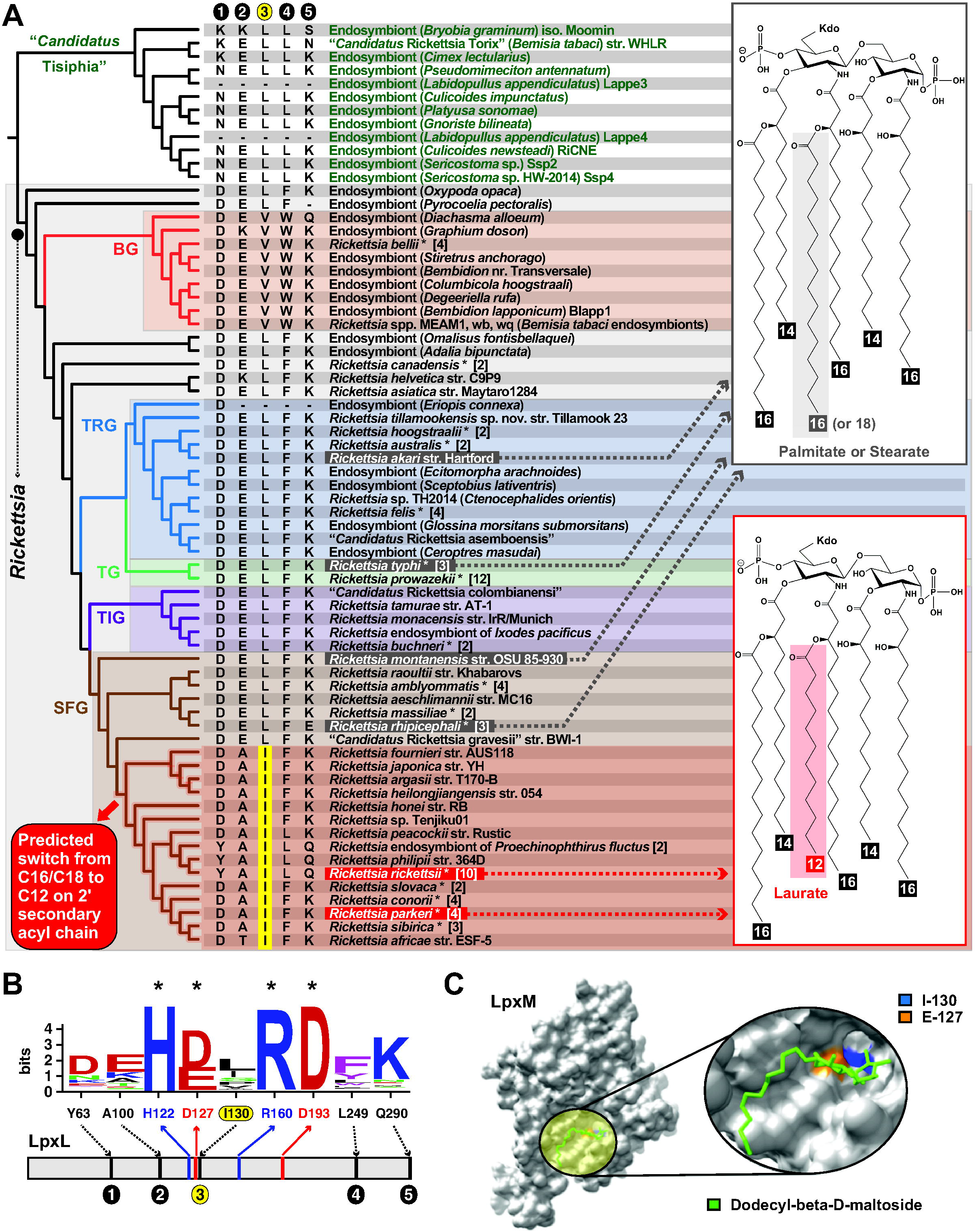
The evolution of variable acyl chain lengths in *Rickettsia* lipid A. **(A)** Superimposition of determined lipid A structures and late acyltransferase LpxL characteristics over an estimated *Rickettsia* phylogeny. Tree is redrawn from a recent study (7). BG, Bellii Group; TRG, Transitional Group; TG, Typhus Group; TIG, Tamurae/Ixodes Group; SFG, Spotted Fever Group. Taxon names in gray boxes were determined to synthesize lipid A with palmitate or stearate (C16 or C18) added to the primary 2’ acyl chain; taxon names in red boxes were determined to synthesize lipid A with palmitate or stearate (C16 or C18) added to the primary 2’ acyl chain (see **Fig. S1** for more details). Red shading indicates the predicted time point in SFG rickettsiae evolution where a switch from palmitate/stearate to laurate on lipid A 2’ hydroxypalmitate could have occurred based on our structural determinations and shared features of LpxL proteins. Circles 1-5 depict the only variable positions from an alignment of LpxL proteins (panel **B**). Yellow highlighting indicates a conserved Ile in variable position 3 for the clade containing species adding laurate on lipid A 2’ hydroxypalmitate. **(B)** Sequence logo (27) showing conservation of acyltransferase active site residues (asterisks) and five variable positions from an alignment of 127 non-redundant rickettsial LpxL proteins. Alignment performed using MUSCLE (default parameters) (28). Complete information for all proteins is provided in **Table S1**. Amino acid coloring scheme and assignment as follows: black, hydrophobic; red, negatively charged; green, hydrophilic; purple, aromatic; blue, positively charged. The five variable positions are shown for taxa in the estimated phylogeny in panel **A. (C)** Visualization of LpxL variable position 3 within the active site of the lipid A late acyltransferase LpxM (Note: LpxM is analogous to *Rickettsia* LpxJ and acylates 3’ acyl chains but was used since no 2’ secondary acyltransferase structures exist). Gray, surface representation of lipid A acyltransferase LpxM from *Acinetobacter baumannii* (PDBID: 5KN7) is shown in grey. Yellow circle highlighting and insert depict n-dodecyl-β-D-maltoside (DDM) ligand in green. The acidic active site position (E-127, orange) has a reduced substrate binding when mutated to Ala. *Rickettsia* variable position 3 (I-130, blue) is found on the same helix near the catalytic E-127 and may be involved in mediating acyl chain length substrate selectivity. Note: 5KN7-N-130 is mutated to I-130 for this depiction; 5KNK is E-127A mutant structure (not shown). Structures visualized using Chimera (29).

Finally, we inspected *Rickettsia* LpxL proteins for possible active site traits that correlate with lipid A structural heterogeneity (**FIG. 2A, B**). LpxL is the late acyltransferase in the Raetz pathway that acylates 2’ primary acyl chains (see **FIG. S1**). Of only five variable residues across *Rickettsia* LpxL proteins, two (positions 100 and 130) have distinct properties defining later-evolving SFG rickettsiae (**FIG. 2A, B**). While nonetheless interesting with short chain (Ala/Thr) replacing charged (Lys/Glu) residues, position 100 is outside of the LpxL active site. However, position 130 (highlighted yellow in **FIG. 2A, B**) has a conserved Iso in place of Val/Leu and is proximal to active site residue Glu-127, which has been implicated in substrate binding for the analogous late acyltransferase LpxM (26) (**FIG. 2C**). This compelling correlation of acyl chain length and active site composition across *Rickettsia* phylogeny warrants characterizing substrate specificities for divergent *Rickettsia* LpxL enzymes.

In sum, our work 1) bolsters *Rickettsia* lipid A structural heterogeneity, 2) shows FLAT^n^ is effective for analyzing obligate intracellular bacterial lipid A, and 3) identifies LpxL properties that may explain variable acyl chain length addition. We anticipate our findings to impact studies comparing host immune responses to divergent pathogens and inform on the efficacy of lipid A in *Rickettsia* vaccinology.

## ACKNOWLEDGMENTS

The authors would like to acknowledge Francesca Gardner, Richard Smith, Matthew Sherman, Jane Michalski, and Alison Scott for their support. We thank Chris Paddock (Center for Disease Control, Atlanta) and Sean Riley (University of Maryland, College Park) for providing isolates of *Rickettsia rhipicephali* and *Rickettsia parkeri*, respectively.

This work was supported with funds from the National Institute of Health/National Institute of Allergy and Infectious Diseases grants (R21AI26108 to JJG, R01AI147314 to RKE, R21AI166832 to JJG and RKE). The content is solely the responsibility of the authors and does not necessarily represent the official views of the funding agencies. The funders had no role in study design, data collection and analysis, decision to publish, or preparation of the manuscript.

## SUPPLEMETAL MATERIAL LEGENDS

**FIGURE S1**. <u>The Raetz pathway for</u> *<u>Rickettsia</u>* <u>lipid A biosynthesis is variable for 2’ secondary acyl chain addition</u>. As *Rickettsia* species lack enzymes for synthesizing amino sugars and pentose phosphates, they are predicted to use host *N*-acetylglucosamine-1-P and ribose-5-P to initiate biosynthesis of lipid IV(A) and 3-deoxy-D-manno-octulosonate (Kdo), respectively (3, 6). The enzymes synthesizing these two molecules (GlmU_N, LpxA, LpxC, LpxD, LpxI, LpxB, LpxK, RpiB, KdsD, KdsA, KdsB, WaaA) are conserved in all *Rickettsia* genomes and ultimately generate Kdo_2_-lipid IV(A). Note: Kdo_2_ is not shown on the final structure as its structure, as well as structures for inner and outer core oligosaccharide, have not been determined for rickettsiae. Variable 2’ secondary acyl chain addition by LpxL is shown at *top-center*, with lauroyl-Kdo_2_-lipid IV(A) synthesized for *R. parkeri* and *R. rickettsii* strains and either palmityl- or stearyl-Kdo_2_-lipid IV(A) synthesized for *R. akari, R. typhi*, and *R. montanensis*. By contrast, 3’ secondary chain acylation is conserved in all generated structures, with palmitic acid added to 3’-hydroxytetradecanoate by the late acyltransferase LpxJ that we previously characterized (30). Enzyme names are accompanied with locus tags for *R. typhi* str. Wilmington.

**TABLE S1**. <u>Rickettsial LpxL sequences used for bioinformatics analyses</u>. *Rickettsia typhi* (AAU04163) was used in a BlastP search against the NCBI ‘Rickettsiales’ database (taxid: 766). Searches were performed with composition-based statistics, with no filter used. Default matrix parameters (BLOSUM62) and gap costs (Existence: 11 Extension: 1) were implemented, with an inclusion threshold of 0.005. All non-redundant subjects (n = 126) were retrieved and used to assess conservation (see **Fig. 2B**). Note: As of May 28^th^, 2023, species in the genus “*Candidatus* Tisiphia” (1) are grouped in NCBI taxid 780 (“Rickettsia”) and were included here as the outgroup to genus *Rickettsia*.

