## Supplementary figures and images for "Structural determination of *Rickettsia* lipid A without chemical extraction confirms shorter acyl chains in later-evolving Spotted Fever Group pathogens"

### Supplemental Figure 1

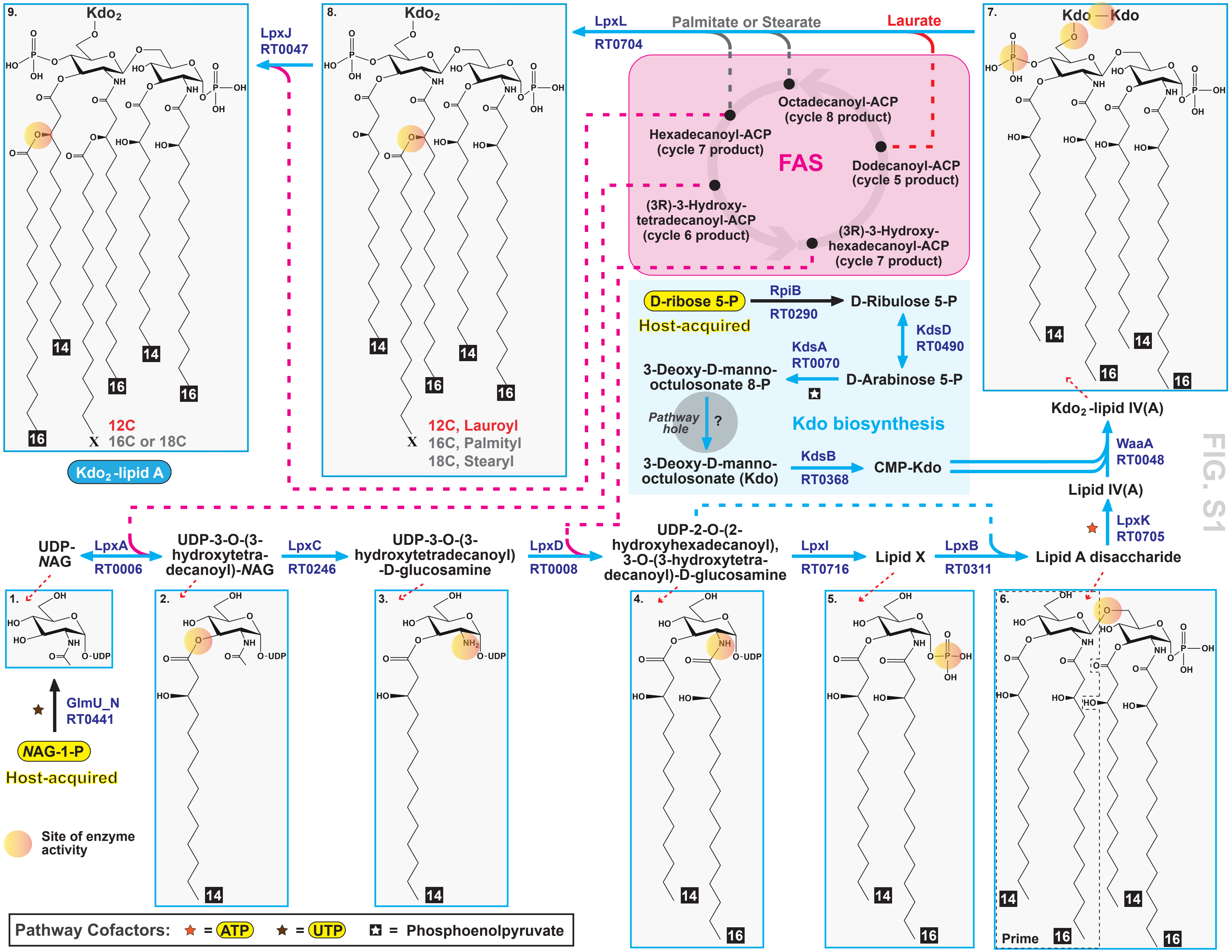
